# Variation in Placental microRNA Expression Associates with Familial Cardiovascular Disease

**DOI:** 10.1101/2021.02.01.429202

**Authors:** Jesse M. Tehrani, Elizabeth M. Kennedy, Fu-Ying Tian, Todd M. Everson, Maya Deyssenroth, Amber Burt, Karen Hermetz, Ke Hao, Jia Chen, Devin C. Koestler, Carmen J. Marsit

## Abstract

In the United States, cardiovascular disease is the leading cause of death, and the rate of maternal mortality remains among the highest of any industrialized nation. Maternal cardiometabolic health throughout gestation and postpartum is representative of placental health and physiology. Both proper placental functionality and placental microRNA expression are essential to successful pregnancy outcomes, and both are highly sensitive to genetic and environmental sources of variation. While placental pathologies, such as preeclampsia, are associated with maternal cardiovascular health and may contribute to the developmental programming of cardiovascular disease, the role of more subtle alterations to placental function and microRNA expression in this relationship remains poorly understood. To develop a more comprehensive understanding of how cardiometabolic health influences placental microRNA expression, and how this shapes placental functionality, we performed small RNA sequencing to investigate microRNA in the placentae from the Rhode Island Child Health Study (n=230). We modeled microRNA counts on maternal family history of cardiovascular disease using negative binomial generalized linear models, and identified microRNAs that were differential expressed (DEmiRs) at a false discovery rate (FDR) less than 0.10. Utilizing parallel mRNA sequencing data and bioinformatic target prediction software, we identified potential mRNA targets of these DEmiRs. We identified 9 DEmiRs, with predicted targets of those miRNA enriched overwhelmingly in the TGFβ signaling pathway but also in pathways involving cellular metabolism and immunomodulation. Overall, we identified a robust association existing between familial cardiovascular disease and placental microRNA expression which may be implicated in both placental insufficiencies and the developmental programming of cardiovascular disease.

## Introduction

As the leading cause of death in the United States, cardiovascular disease (CVD) is expected to affect 44.1% of the population by 2035 ^[1]^. CVD has an extremely complex, multifactorial etiology, where neither genetic nor environmental influences act alone to contribute to disease onset ^[2, 3]^. There is growing body of literature linking maternal cardiometabolic health during pregnancy to adverse cardiovascular health outcomes in offspring later in life ^[4, 5]^. These developmental origins of long-term cardiovascular health outcomes are thought to be the consequence of changes to placental physiology.

The state of a mother’s cardiometabolic health during gestation and postpartum is related to the health and functionality of the placenta. As a central vascular organ overseeing fetal growth, development and the intrauterine environment, proper functionality of the placental remains central to successful gestational outcomes. Resting at the interface of the maternal and fetal environment, the placenta participates in a variety of molecular processes, such as nutrient transport, immunomodulation, and endocrine signaling, all of which are made possible through the development of a villi-based vasculature system, allowing for communication between mother and offspring ^[6]^. Maternal cardiovascular and metabolic conditions during pregnancy, such as type II diabetes, hypertension and hypercholesterolemia have been implicated in dysfunction of placental micro and macro-vasculature. In addition, pathophysiologic characteristics of placental insufficiency, such as those occurring in preeclampsia, manifests in cardiovascular effects including hypertension and kidney malfunction ^[7, 8]^. Additionally, such placental insufficiencies are associated with adverse gestational outcomes, such as fetal growth restriction, which itself serves as a significant risk indicator for the development of cardiovascular disease (CVD) later in life ^[9-11]^. Considering placental development occurs concurrently with fetal heart development, and each organ utilizes common humoral growth signals, such as transforming growth factor-beta (TGFβ) and vascular endothelial growth factor (VEGF), deficiencies in the organogenesis of either organ may alter the formation of the other, initiating physiological changes with potential lifelong cardiovascular consequences ^[12-14]^. While the associations between maternal cardiometabolic risk factors, placental insufficiency and offspring lifelong health outcomes are well defined ^[7, 8, 15]^, the molecular mechanisms by which this developmental programming is established have yet to be robustly delineated. Additionally, these overt placental pathologies are relatively rare, and cannot account for the prevalence of cardiovascular disease and deficiencies in maternal health, post-partum.

MicroRNAs (miRNA) are small noncoding RNA molecules (∼22 nucleotides) capable of post-transcriptional regulation of gene expression. These molecules utilize base-pairing to bind to the 3’-untranslated region of target mRNAs resulting in either translational repression or mRNA degradation, by which the exact mechanism largely depends on the degree of sequence complementarity between the miRNA and target mRNA. Dysregulation of placental miRNA have previously been implicated in both preeclampsia and fetal growth restriction, suggesting their potential role as modifiers of newborn and maternal health outcomes in response to various genetic and/or environmental conditions ^[16-21]^. However, many of these studies were conducted with on small sample sizes (n<100), and largely focused on newborn outcomes as they associate with placental miRNA expression as opposed to the relationship between maternal characteristics and expression of placental miRNA ^[16-19, 21]^.

To explore the influence of familial cardiovascular disease on the placental miRNA landscape, we utilized placental miRNA sequencing data from the Rhode Island Child Health Study (RICHS; n=230) and examined the relationship between maternal family history of CVD and placental miRNA expression. Through this clinical history variable, we capture both genetic risk of CVD, as well as shared familial behaviors that may associate with CVD, such as diet and exercise ^[3, 22]^. Bioinformatic target prediction was used to identify potential mRNA targets of miRNAs significantly associated with maternal CVD risk, followed by overrepresentation analyses to characterize the biochemical pathways in which these mRNAs participate.

## Results

This study analyzed miRNA sequencing data from 230 placentae from the Rhode Island Child Health Study (RICHS). The demographics of the participants are displayed in Table 1. In general, placentae collected in this study were from full term pregnancies (≥37 weeks), all from relatively healthy mothers who did not experience serious pregnancy complications. Forty-nine percent (n=113 of 230) of maternal participants reported a family history of cardiovascular disease, and forty-nine percent (n=112 of 230) were either overweight or obese, n=57 of those also reporting a family history of CVD.

**Table 1.** Demographic characteristics of participants included in the miRNA sequencing analysis (n=230)

| **MATERNAL CHARACTERISTICS** |  |
| --- | --- |
| **Age (yrs) – mean (range)** | 30.9 (18 - 40) |
| **Family History of CVD (Yes) – % (n)** | 49% (112) |
| **Race (White) – % (n)** | 77% (178) |
| **Pregnancy Smoking – % (n)** | 9% (20) |
| **Pre-Pregnancy Body Mass Index (BMI) (kg/m<sup>2</sup>) – mean (range)** | 26.5 (16 - 46.7) |
| **Pre-Pregnancy BMI Group** |  |
| Normal and Underweight – % (n) | 51.3% (118) |
| Overweight – % (n) | 27.0% (62) |
| Obese – % (n) | 21.7% (50) |
| **INFANT CHARACTERISTICS** |  |
| **Gestational Age (wks) – mean (range)** | 39.4 (37 - 41.4) |
| **Sex (Female) – % (n)** | 49% (113) |
| **Birthweight (g) – mean (range)** | 3563.7 (2030 - 5465) |
| **Birthweight Group** |  |
| Small for Gestational Age (SGA) – % (n) | 14.3% (33) |
| Average for Gestational Age (AGA) – % (n) | 54.3% (125) |
| Large for Gestational Age (LGA) – % (n) | 31.3% (72) |

To analyze the association between family history of CVD and placental miRNA expression, we performed differential expression analysis using negative binomial generalized linear models constructed in *DESeq2* on placental small RNA sequencing data. We identified 9 <u>**D**</u>ifferentially <u>**E**</u>xpressed <u>**miR**</u>NAs (DEmiRs) associated with familial CVD history (FDR < 0.1), 4 of which (miR-1246, miR-324-5p, miR-1307-3p, and miR-520a-3p) met a strict Bonferroni threshold (p-value < 6.23e-05) (Fig. 1). Sensitivity analyses were performed to adjust for various biological and technical covariates (RNA integrity, flow cell lane, pregnancy smoking, maternal race, infant sex and infant birth weight percentile) in order to characterize the robustness of DEmiR effect sizes, and the DEmiR log_2_ fold changes were striking consistent even with these additional adjustments to the original model (Fig. S1 & Fig. S2).

**Figure 1:**
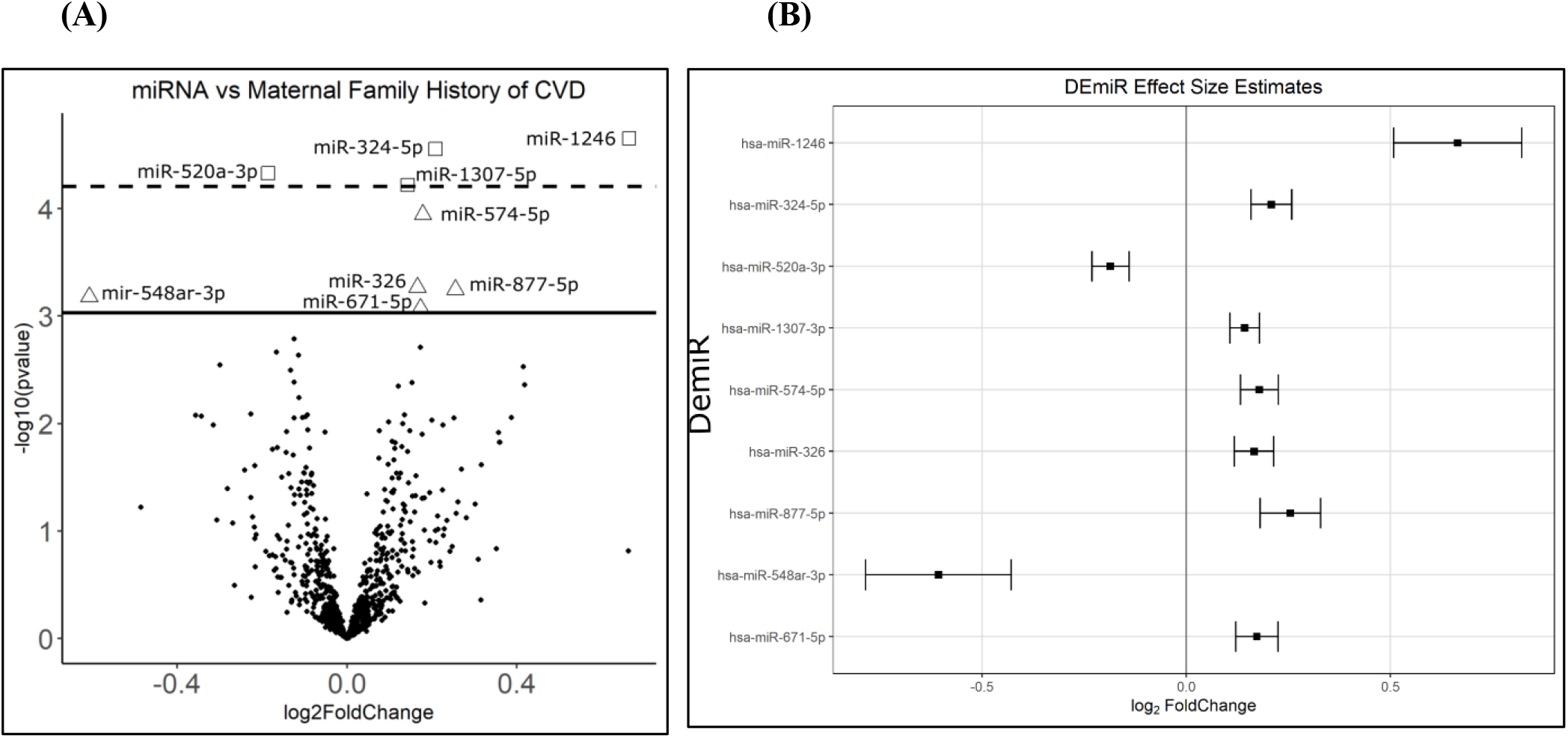
Placental miRNA associates with maternal family history of CVD. Volcano plot representing the results of the differential expression analysis. The y-axis shows the -log_10_(p-values) in the association of each miRNA with family history of CVD. The x-axis displays the effect estimates in units of log_2_ fold change in each miRNA’s transcript abundance in individuals with familial incidence of CVD. 9 miRNAs are significantly (FDR <0.1) associated with familial CVD history and 4 of those (square shaped) reach are significant after Bonferroni correction (p-value < 6.23e-05). **(B)** Estimates of log_2_ fold change of miRNA transcript abundance of miRNAs significantly associated with familial CVD history. Boxes represent the effect size estimate, while error bars represent standard error of the effect size estimate.

Bioinformatic targets of all 9 DEmiRs were predicted using the miRNA Data Integration Portal (miRDIP)^[23]^. miRDIP predicted 4,516 targets across all 9 DEmiRs. However, given that miRNA do not require perfect complementarity with a target mRNA and can only target mRNAs expressed in the examined tissue, bioinformatic target prediction is prone to generating false positives. To enhance our dataset with true miRNA:mRNA relationships we utilized parallel total RNA sequencing data in RICHS ^[24]^. Only miRNA:mRNA pairs where the transcript abundance of a DEmiR and its miRDIP predicted target were negatively correlated (Pearson’s correlation, p-value < 0.05) were considered in downstream pathway analyses. This correlation-based filtering revealed 7 DEmiRs and 66 predicted mRNA targets (Table S1) ^[24]^.

The 66 predicted DEmiR targets were tested for pathway overrepresentation within ConsensusPathDB (CPDB)^[24-26]^. Pathways related to TGFβ signaling, cellular metabolism, and immunomodulation were overrepresented among DEmiR targets (q-value < 0.05). These enrichment results are largely driven by targets of miR-574-5p, miR-324-5p, miR-326 and miR-520a-3p (Table S1).

## Discussion

Maternal mortality rates in the United States are higher than any other industrialized country and continue to rise. More than half of these deaths are deemed preventable, and over 15% of these deaths are caused by CVD ^[27]^. While our study does not investigate placentae from mothers diagnosed with CVD, the chronic nature of the disease provides a unique opportunity to study how minor insults and risk factors, such as family history, influence placental functionality and their contributions to adverse newborn health outcomes.

Our small RNA-seq analysis identified fetoplacental miRNAs whose expression associate with maternal family history of CVD (Fig. 1). The functional relevance of miRNAs is largely dictated by the mRNAs available for them to interact with. These interactions are heavily influenced by the tissue of origin of both miRNAs and their target mRNAs. While many of the DEmiRs we identified (miR 324-5p, miR-520a-3p, miR-574-5p, miR-326) are implicated in cardiometabolic conditions during pregnancy, including preeclampsia and diabetes ^[28-31]^, their functional role remains largely understudied in the context of placental physiology.

Pathway overrepresentation analyses of DEmiR predicted targets overwhelmingly suggested TGFβ signaling to be influenced by dysregulation of placental miRNAs whose expression associates with family history of CVD (Table 2). TGFβ signaling encompasses a large family of genes that participate in a myriad of cellular processes, but these genes are particularly important to vascular tissues for neovascularization as well as remodeling and repair of existing tissue ^[32]^. Among our predicted DEmiR targets, specifically those of miR-574-5p, are SMAD2 and SMAD4, two activators of the TGFβ pathway whose dysregulation are implicated in preeclampsia pathogenesis in human umbilical vein cells (HUVEC) *in vitro* ^[33-35]^. SMAD2/4 also appear frequently throughout the various biological pathways reported as overrepresented, many of which contribute toward TGFβ signaling (including activin and nodal signaling pathways), highlighting their importance to a diverse set of endothelial cellular processes and placental physiology as a whole (Table 2) ^[36]^. In addition, while miR-1246 targets did not pass our strict filtering criteria to be included in the pathway analysis, upregulation of miR-1246 has been shown to increase angiogenesis through activation of TGFβ signaling in HUVEC ^[37]^.

**Table 2.** DEmiR Target Pathway Enrichment Analysis Results.

| **Pathway** | **Enriched Pathway Components** | **p-Value** | **q-Value** | **Source** |
| --- | --- | --- | --- | --- |
| **Signaling by Activin** | SMAD4; ACVR2B; SMAD2 | 2.5E-05 | 0.00196 | Reactome |
| **Signaling by NODAL** | SMAD4; ACVR2B; SMAD2 | 2.5E-05 | 0.00196 | Reactome |
| **Activation of BAD and translocation to mitochondria** | AKT3; YWHAQ; BCL2 | 6.8E-05 | 0.00278 | Reactome |
| **Intrinsic Pathway for Apoptosis** | AKT3; BCL2L1; YWHAQ; BCL2 | 7.7E-05 | 0.00278 | Reactome |
| **The NLRP1 inflammasome** | BCL2L1; BCL2 | 8.8E-05 | 0.00278 | Reactome |
| **Role of Calcineurin-dependent NFAT signaling in lymphocytes** | RAN; BCL2L1; YWHAQ; BCL2 | 1.7E-04 | 0.00435 | PID |
| **Downregulation of SMAD2/3:SMAD4 transcriptional activity** | SMAD4; NEDD4L; SMAD2 | 1.9E-04 | 0.00435 | Reactome |
| **Platelet degranulation** | APP; APLP2; PHACTR2; TAGLN2; CALU | 3.4E-04 | 0.00665 | Reactome |
| **Response to elevated platelet cytosolic Ca<sup>2+</sup>** | APP; APLP2; PHACTR2; TAGLN2; CALU | 4.0E-04 | 0.00697 | Reactome |
| **Activation of BH3-only proteins** | AKT3; YWHAQ; BCL2 | 5.1E-04 | 0.00812 | Reactome |
| **TGF-beta receptor signaling activates SMADs** | SMAD4; NEDD4L; SMAD2 | 6.9E-04 | 0.00990 | Reactome |
| **BH3-only proteins associate with and inactivate anti-apoptotic BCL-2 members** | BCL2L1; BCL2 | 8.1E-04 | 0.01064 | Reactome |
| **Class I PI3K signaling events mediated by Akt** | AKT3; BCL2L1; YWHAQ | 9.0E-04 | 0.01093 | PID |
| **TGF-Core** | SMAD4; ACVR2B; SMAD2 | 9.8E-04 | 0.01103 | Signalink |

TGFβ signaling in the placenta is known to play critical roles in preimplantation and initial decidualization, and is also active throughout gestation in endometrial remodeling ^[33, 34]^. While neovascularization is required for successful fetal development, aberrant regulation of the signaling pathways overseeing this process may lead to improper vessel network development in the form hypervascularization or vasculature malformation ^[38, 39]^. Additionally, these structural abnormalities are associated with placental insufficiencies, which could result in lifelong adverse cardiometabolic health outcomes in both mothers and offspring ^[12, 13, 40]^. While the data presented here are incapable of concluding the direct impact of placental miRNA dysregulation there is a growing body of literature, as well as the associations outlined here, that suggest dysregulation of placental miRNAs may be contributing to the developmental programming and propagation of CVD. However, these findings should be interpreted within the context of this study’s limitations.

This study only includes term placentae from live births, where premature births and other birth defects were excluded. The cross-sectional design of this study limits the interpretation of miRNA associations temporally, and may not be representative of miRNA associations throughout gestational development. Additionally, this study relies on self-reported family history of CVD, and which may ultimately lead to variation in results attributable to recall bias.

## Conclusion

MiRNAs serve as an important form of post-transcriptional gene regulation during early development, and are sensitive to both genetic and environmental conditions. Here we have shown that the expression of 9 placental miRNAs are associated with maternal family history of CVD, and that the mRNA targets of these miRNAs largely play a role in TGFβ signaling, indicating their involvement in endothelial cell functionality and placental physiology as a whole. Dysregulation of miRNA expression in the placenta may contribute to adverse newborn and maternal health outcomes, ultimately playing a role in the developmental programming of CVD.

## Methods

### Cohort

***The Rhode Island Child Health Study (RICHS)*** is a cohort of mother-infant pairs from the Women & Infants Hospital in Providence, Rhode Island, enrolled between September 2010 and February 2013. All mothers were at least 18 years of age, had no life-threatening conditions, and delivered singletons free of congenital/chromosomal abnormalities at or after 37 weeks of gestation. All participants provided written informed consent and all protocols were approved by the IRBs at the Women & Infants Hospital of Rhode Island and Emory University, respectively. Data provided by this study include placental microRNA transcript abundance (n=230). Interviewer-administered questionnaires were utilized to collect sociodemographic and lifestyle data. Structured medical record review was used to collect anthropometric and medical history data. Maternal family history of CVD was reported and coded as a binary variable (yes/no). Follow-up questions regarding which primary relative had been afflicted, coded as “mother”,” father”,” brother”, and “sister”. Instances where the mother reported a family history of CVD but did not identify a specific primary relative were not included in the study in order to prevent improper classification of familial CVD incidence.

### Tissue Collection

Fetal placental samples from all subjects were collected as previously described^[41]^. Briefly, placental samples were collected within two hours of birth; fragments were obtained two centimeters (cm) from the umbilical cord and free of maternal decidua. Collected tissue was immediately placed in RNA later solution (Life Technologies, Grand Island, NY, USA) and stored at 4 °C for at least 72 hours. Subsequently, tissue segments were blotted dry, snap frozen in liquid nitrogen, homogenized by pulverization using a stainless-steel cup and piston unit (Cellcrusher, Cork, Ireland) and stored at −80 °C.

### miRNA isolation and sequencing

Total RNA was extracted from placenta using the Qiagen miRNeasy Mini Kit and a TissueLyser LT (Qiagen, Germantown, MD, USA) following manufacturer’s protocol. Briefly, 25-35 mg of frozen, powdered placental tissue was placed in a 2 ml round bottom tube with 700 ul of Qiazol Lysing Reagent and one 5 mm stainless steel bead. The tissue was homogenized in a pre-chilled tube holder on the TissueLyser LT for two, 5-minute cycles at 30 Hz. The resulting homogenate was processed with the Qiagen miRNeasy Mini Kit with on-column DNAse digestion and eluted in 50 µl RNase-free water. The RNA was quantitated on a NanoDrop (Thermo Fisher, Waltham, MA, USA) and quality checked on Agilent Bioanalyzer using the Agilent RNA 6000 Nano kit (Agilent, Santa Clara, CA, USA). Single end, 1 x 50 bp next generation sequencing of placental miRNA was performed by Omega Bioservices (Norcross, Georgia) as previously described ^[24]^.

### miRNA Seq Processing and QC

Raw FASTQ reads obtained from a total of 230 RICHS samples were subject to adaptor trimming with cutadapt v1.1634. The 3’ adaptor sequence were trimmed (TGGAATTCTCGGGTGCCAAGG) and then four bases were trimmed from each end of the read following vendor’s recommendation (BIOO scientific, Austin TX). We then used trimmed reads and miRDeep2 to quantify microRNA ^[42]^. miRDeep2 was used to first perform alignment using bowtie1 with human genome hg38 ^[43]^. The ‘Quantifier’ module in miRDeep2 was used to obtain raw counts of microRNAs with miRBase version 22 ^[44]^.

### Transcript Filtering and Normalization

Raw miRNA counts were imported into *DESeq2* for normalization and differential expression analysis. Only miRNA transcripts with more than one count per million in at least 10 percent of samples were included, leaving 802 miRNA transcripts to be analyzed of the initial 2656 sequenced transcripts. Dispersion estimates were then calculated, followed by generation of median ratio size factor estimates to normalize counts for analysis with in *DESeq2* ^[45]^. Normalized counts were then exported from *DESeq2* for Surrogate Variable Analysis (SVA). The Variance Stabilization Transformation (VST) was applied to count matrices to yield approximately normalized and log_2_-transformed abundances, which were utilized in correlation analyses ^[24]^.

## Statistical Analyses

### SVA

In an effort to adjust for unknown confounders, such as cell-type heterogeneity and unmeasured sources of technical variation, surrogate variables were estimated for normalized miRNA transcript reads using the *sva* package ^[46, 47]^. The full model used in the *svaseq* includes variation attributable to family history of CVD while the null model included only an intercept term. One surrogate variable was utilized as a covariate in our differential expression analysis.

### DESeq2 Differential Expression Analysis

miRNA transcript counts were modeled using a negative binomial generalized linear model to identify differentially expressed transcripts in *DESeq2* ^[47]^. For each of the 802 individual miRNA transcripts which passed strict filtering and quality criteria, the variance stabilized transformed transcript abundance was regressed on the history of familial CVD (No = 0, Yes = 1), while adjusting for the first surrogate variable to control for unknown confounders^[46, 47]^. We considered miRNAs with a false discovery rate (FDR) less than 10% to be considered a differentially expressed miRNA (DEmiR) with respect to family history of CVD. Extensive sensitivity analyses were performed by including various parameters as covariates to the original model to assess the robustness DEmiR effect sizes (Fig. S1).

### Target Prediction and Filtering

Potential DEmiR targets were identified using miRDIP, an online database of miRNA target predictions ^[23]^. Only targets within the top 1% of confidence scores were returned. We then calculated Pearson correlation coefficients for each miRNA:mRNA pair utilizing normalized miRNA and mRNA sequencing counts from RICHS^[24]^. Only miRNA:mRNA target pairs with negative correlation coefficients (q-value < 0.05) were returned to be used in network and pathway analyses. Correlation coefficients ranged between -0.41 and -0.21.

### Pathway Analysis

Predicted DEmiR targets were tested for pathway overrepresentation within ConsensusPathDB (CPDB) ^[25, 26]^, against all genes that passed general QC filtering in RICHS whole transcriptome RNA-seq analysis ^[24-26]^. CPDB utilizes 12 separate biological pathway databases, and calculates an enrichment p-value from the hypergeometric distribution of genes in the list of miRNA targets and the pathway gene set. Only mRNAs that were expressed >1cpm in at least 10% of the RICHS samples were included as the background for the pathway analysis ^[24]^. False discovery rates were calculated from the enrichment p-values, and a q-value less than 0.05 was considered a significant enrichment of miRNA targets in the tested pathway.

## Supplemental Figures and Tables

**Fig S1:**
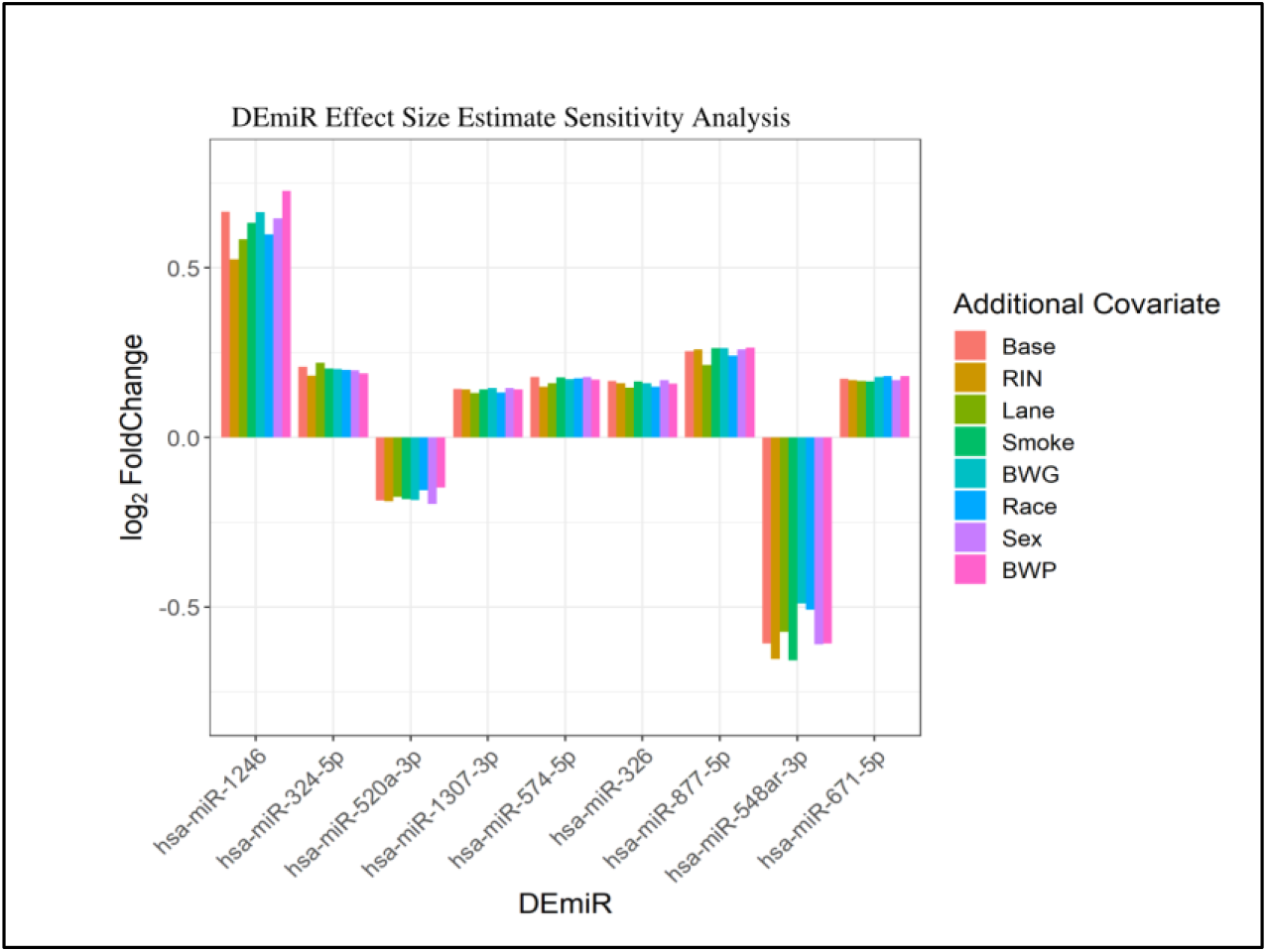
Effect size estimates of DEmiRs are highly robust to various suspected technical and biological covariates. Bar plot showing effect size estimates for all DEmiRs after correction for suspected technical and biological covariates not included in the original (base) model. RIN = RNA integrity, Lane = Flow cell lane, Smoke = maternal smoking status, BWG = birth weigh group, Race = maternal race, Sex = offspring sex, YOB = year of birth, BWP = birth weight percentile.

**Fig S2:**
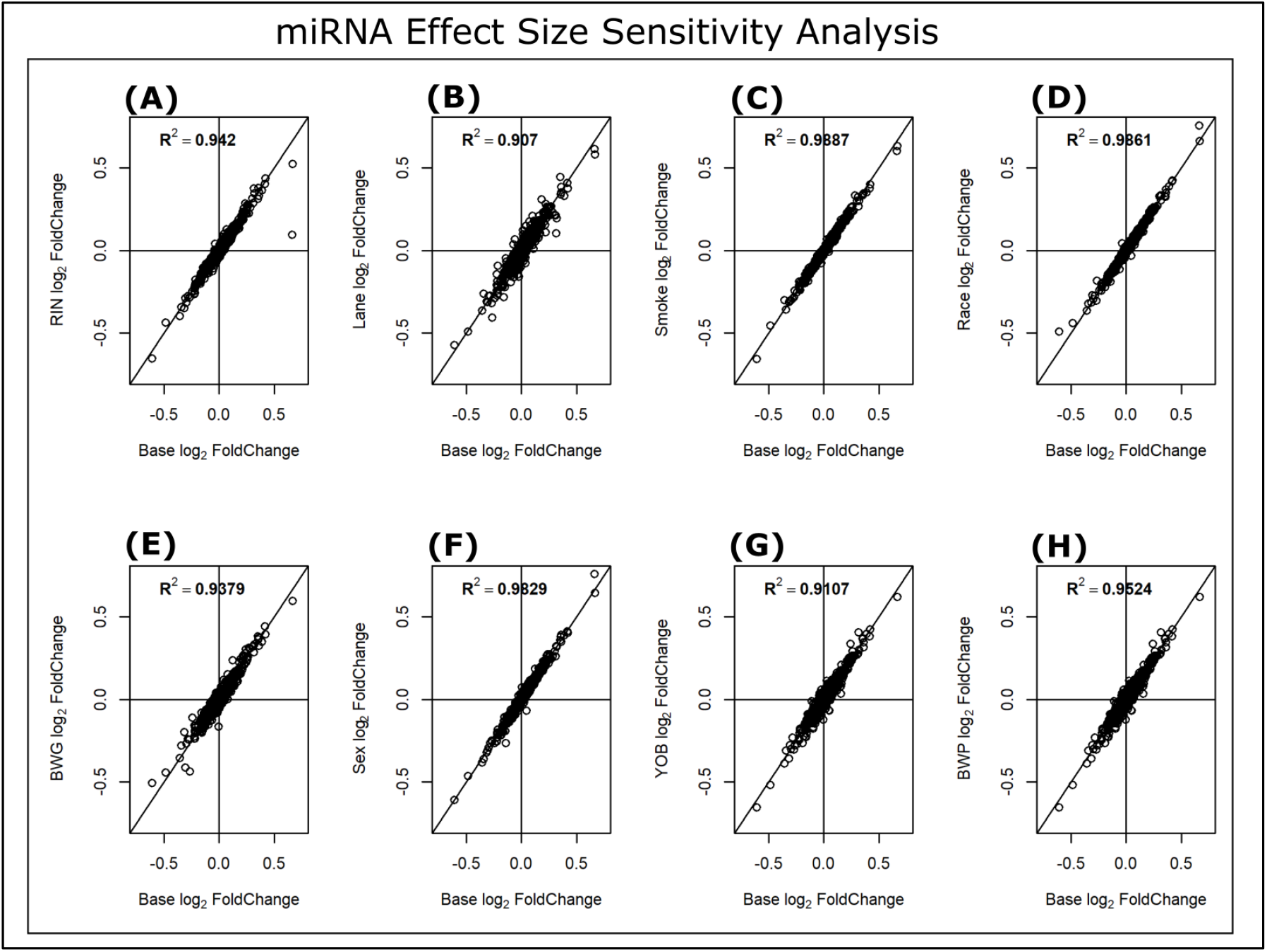
Effect size estimates of placental miRNAs are highly robust on a transcriptome-wide scale to various suspected technical and biological covariates. Scatter plots displaying the coefficient of determination (R^2^) to emphasize the relationship between the estimates produced by the original (base) model and estimates of various models which correct for additional, suspected technical and biological covariates across all miRNAs analyzed. **(A)** RIN = RNA integrity, **(B)** Lane = Flow cell lane, **(C)** Smoke = maternal smoking status, **(D)** Race = maternal race, **(E)** BWG = birth weight group, **(F)** Sex = offspring sex, **(G)** YOB = year of birth, **(H)** BWP = birth weight percentile.

**Table S1.** Filtered miRDIP Output: Predicted DEmiR mRNA Targets

| MicroRNA | Gene Symbol | Uniprot ID | Integrated Score |
| --- | --- | --- | --- |
| hsa-miR-324-5p | VDAC1 | P21796 | 0.81 |
| hsa-miR-324-5p | APP | P05067 | 0.73 |
| hsa-miR-324-5p | RAN | P62826 | 0.73 |
| hsa-miR-324-5p | KLF3 | P57682 | 0.72 |
| hsa-miR-324-5p | KCTD20 | Q7Z5Y7 | 0.63 |
| hsa-miR-324-5p | RACGAP1 | Q9H0H5 | 0.54 |
| hsa-miR-324-5p | DRAM1 | Q8N682 | 0.54 |
| hsa-miR-324-5p | SLC16A1 | P53985 | 0.53 |
| hsa-miR-324-5p | YWHAQ | P27348 | 0.49 |
| hsa-miR-324-5p | ST8SIA4 | Q92187 | 0.46 |
| hsa-miR-324-5p | H2AFV | Q71UI9 | 0.45 |
| hsa-miR-324-5p | TRIP13 | Q15645 | 0.42 |
| hsa-miR-324-5p | CCT5 | P48643 | 0.39 |
| hsa-miR-324-5p | APLP2 | Q06481 | 0.39 |
| hsa-miR-326 | BSDC1 | Q9NW68 | 0.68 |
| hsa-miR-326 | PPP4R1 | Q8TF05 | 0.63 |
| hsa-miR-326 | BCL2L1 | Q07817 | 0.52 |
| hsa-miR-326 | MAD2L1BP | Q15013 | 0.51 |
| hsa-miR-326 | TES | Q9UGI8 | 0.50 |
| hsa-miR-326 | NSUN2 | Q08J23 | 0.47 |
| hsa-miR-326 | JOSD1 | Q15040 | 0.45 |
| hsa-miR-326 | APMAP | Q9HDC9 | 0.43 |
| hsa-miR-326 | RHOC | P08134 | 0.41 |
| hsa-miR-326 | EIF4A1 | P60842 | 0.40 |
| hsa-miR-326 | TAGLN2 | P37802 | 0.39 |
| hsa-miR-520a-3p | FEM1C | Q96JP0 | 0.77 |
| hsa-miR-520a-3p | PKD2 | Q13563 | 0.75 |
| hsa-miR-520a-3p | TRPV6 | Q9H1D0 | 0.73 |
| hsa-miR-520a-3p | SLC25A27 | O95847 | 0.70 |
| hsa-miR-520a-3p | ZNF697 | Q5TEC3 | 0.63 |
| hsa-miR-520a-3p | GLUL | P15104 | 0.56 |
| hsa-miR-520a-3p | NEDD4L | Q96PU5 | 0.52 |
| hsa-miR-520a-3p | SH3TC2 | Q8TF17 | 0.44 |
| hsa-miR-520a-3p | RNASEL | Q05823 | 0.42 |
| hsa-miR-548ar-3p | CLASP2 | O75122 | 0.40 |
| hsa-miR-574-5p | MKRN1 | Q9UHC7 | 0.61 |
| hsa-miR-574-5p | G3BP2 | Q9UN86 | 0.59 |
| hsa-miR-574-5p | BDNF | P23560 | 0.59 |
| hsa-miR-574-5p | PAPPA | Q13219 | 0.55 |
| hsa-miR-574-5p | SEL1L | Q9UBV2 | 0.52 |
| hsa-miR-574-5p | AKT3 | Q9Y243 | 0.50 |
| hsa-miR-574-5p | KLHL18 | O94889 | 0.50 |
| hsa-miR-574-5p | PAK3 | O75914 | 0.49 |
| hsa-miR-574-5p | RAB3B | P20337 | 0.46 |
| hsa-miR-574-5p | CEP350 | Q5VT06 | 0.46 |
| hsa-miR-574-5p | SMAD4 | Q13485 | 0.45 |
| hsa-miR-574-5p | SMAD2 | Q15796 | 0.45 |
| hsa-miR-574-5p | GPR107 | Q5VW38 | 0.44 |
| hsa-miR-574-5p | OSBPL2 | Q9H1P3 | 0.43 |
| hsa-miR-574-5p | CALU | O43852 | 0.42 |
| hsa-miR-574-5p | ZCCHC2 | Q9C0B9 | 0.41 |
| hsa-miR-574-5p | MAN1C1 | Q9NR34 | 0.41 |
| hsa-miR-574-5p | ACVR2B | Q13705 | 0.41 |
| hsa-miR-574-5p | PCM1 | Q15154 | 0.41 |
| hsa-miR-574-5p | PHACTR2 | O75167 | 0.40 |
| hsa-miR-574-5p | BCL2 | P10415 | 0.40 |
| hsa-miR-574-5p | KCMF1 | Q9P0J7 | 0.39 |
| hsa-miR-574-5p | HS3ST3B1 | Q9Y662 | 0.39 |
| hsa-miR-574-5p | IL1RAP | Q9NPH3 | 0.39 |
| hsa-miR-574-5p | CANX | P27824 | 0.39 |
| hsa-miR-574-5p | ARFGEF2 | Q9Y6D5 | 0.39 |
| hsa-miR-574-5p | NAB2 | Q15742 | 0.39 |
| hsa-miR-574-5p | ASNA1 | O43681 | 0.38 |
| hsa-miR-671-5p | VPS26A | O75436 | 0.43 |
| hsa-miR-877-5p | REV3L | O60673 | 0.56 |

**Table S2:**
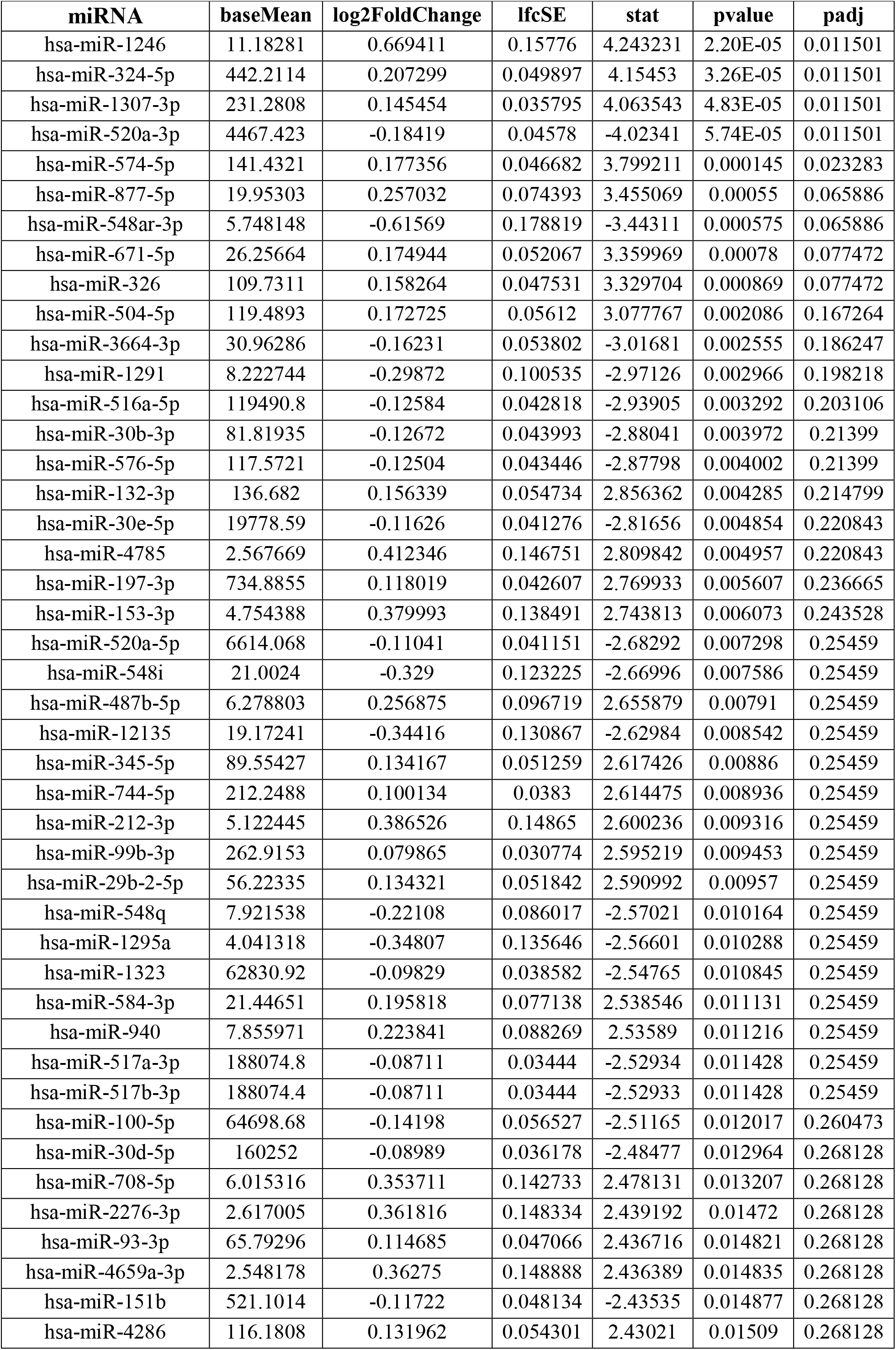
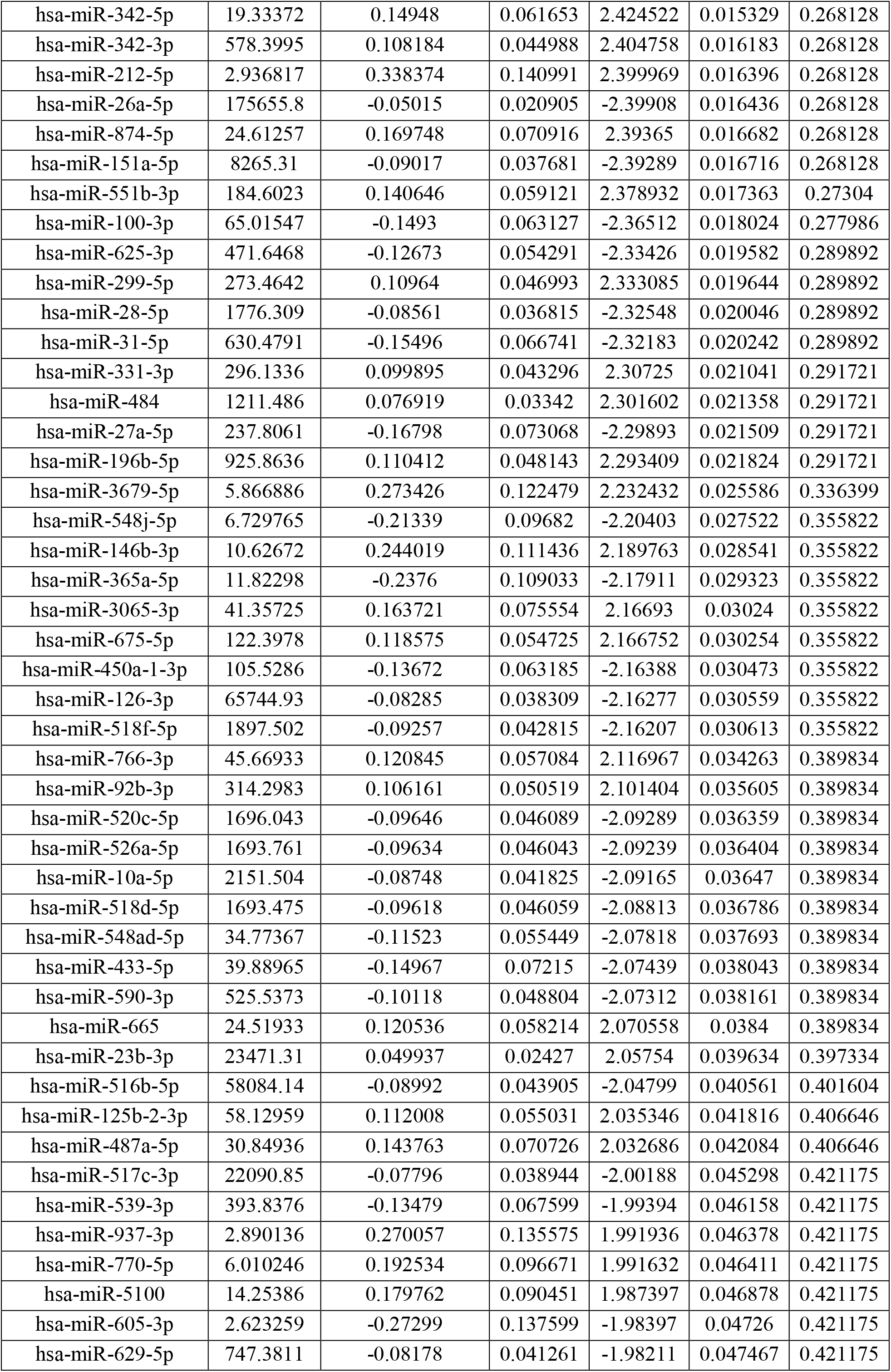
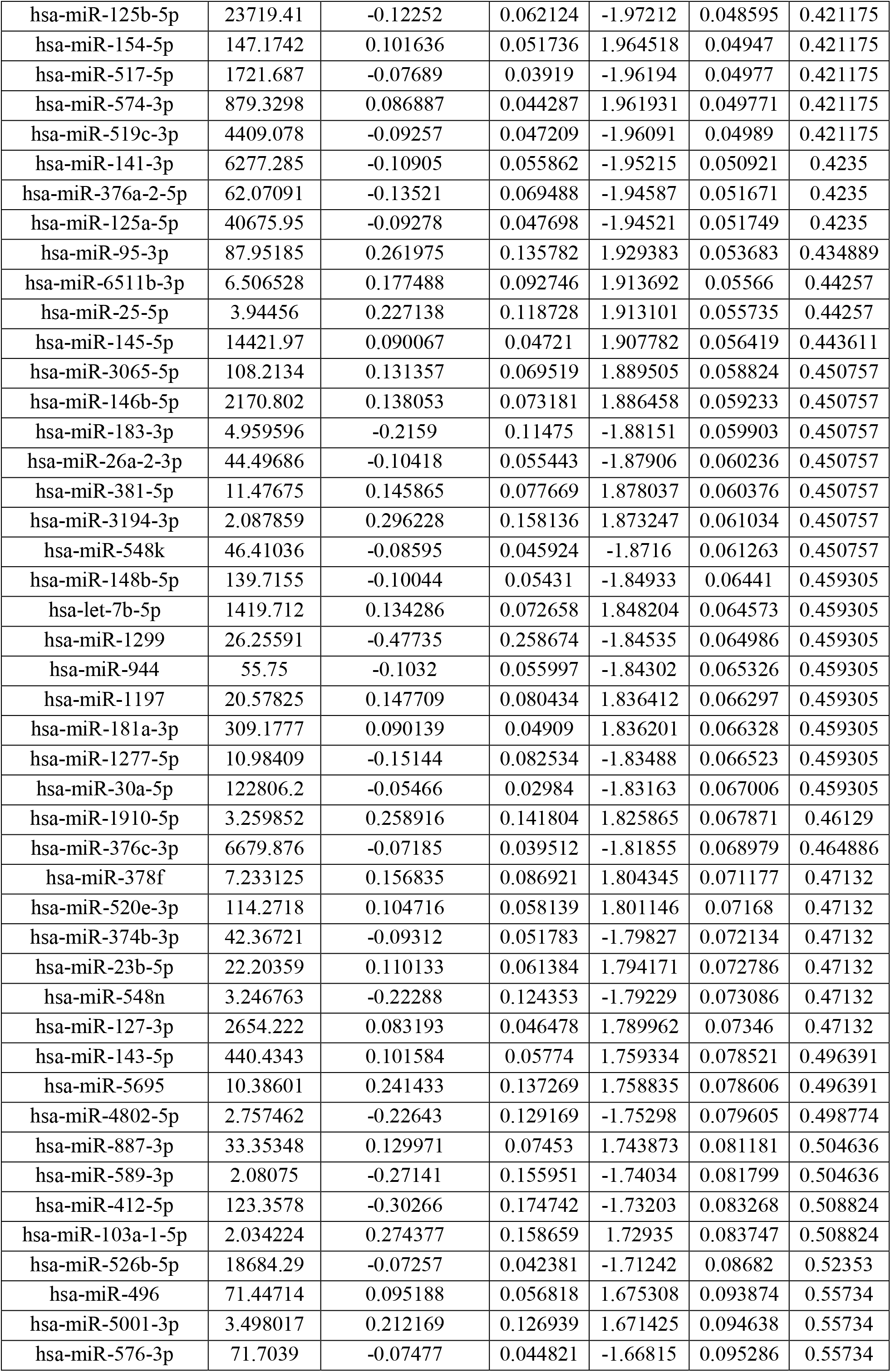
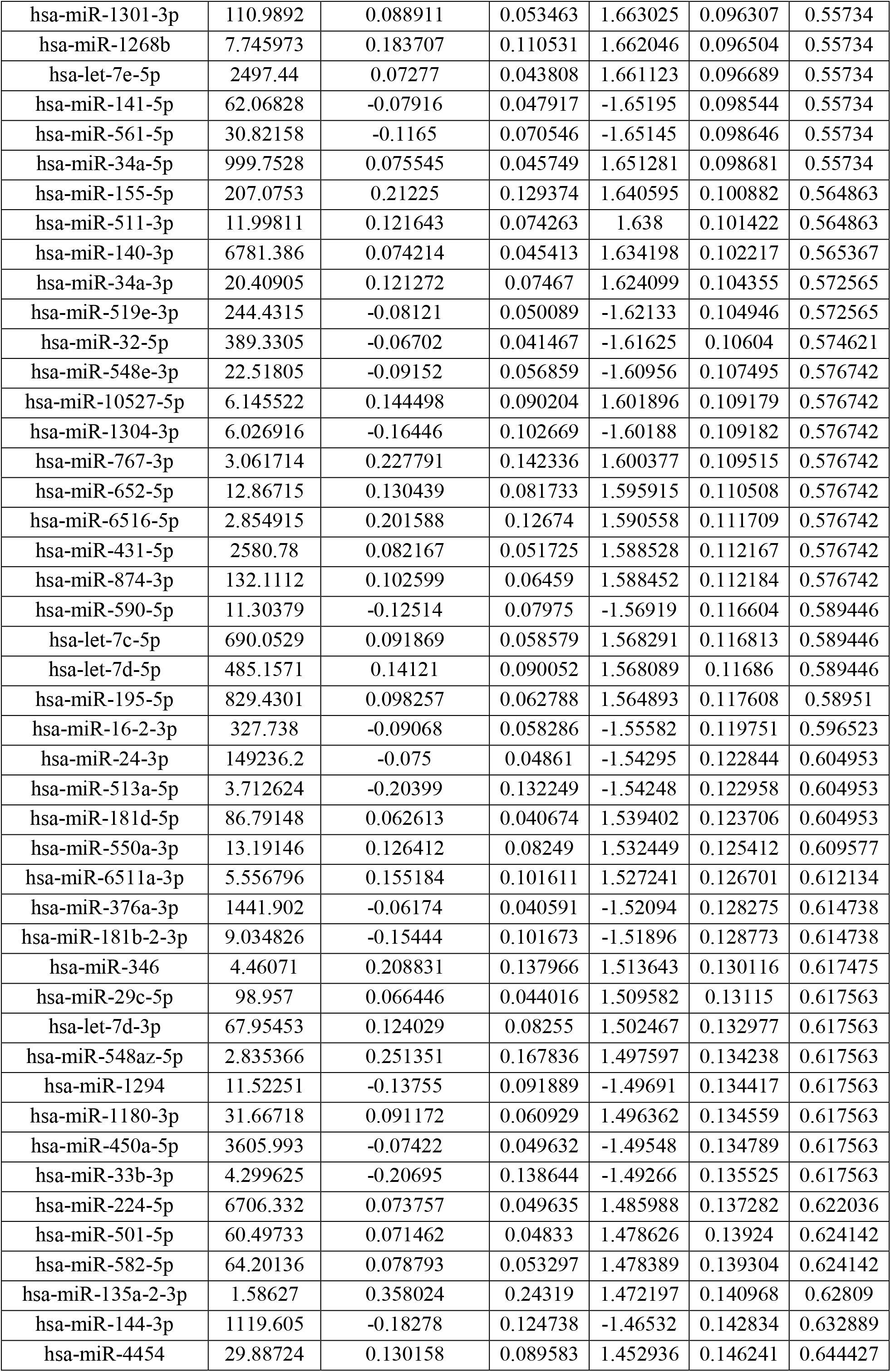
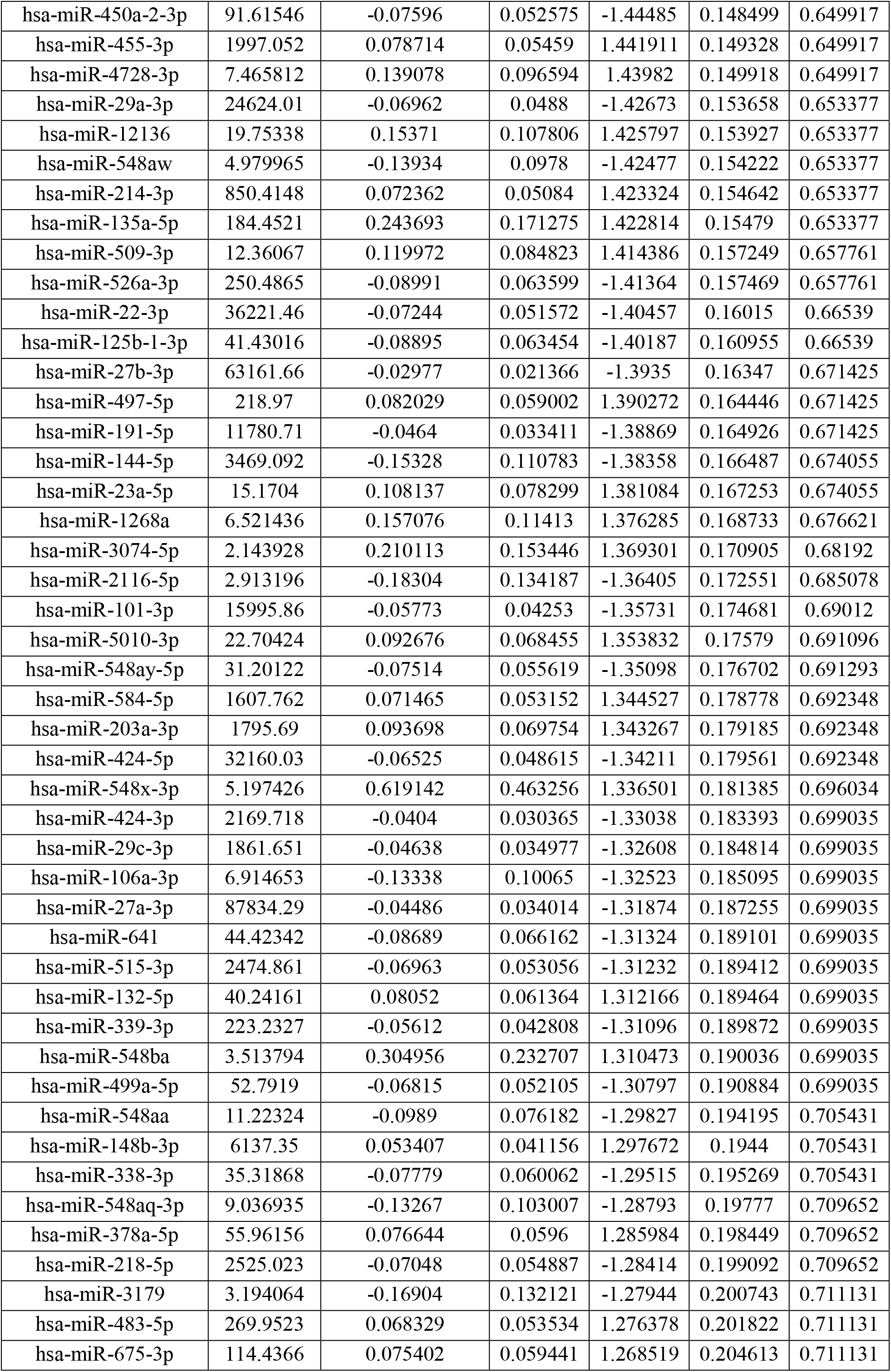
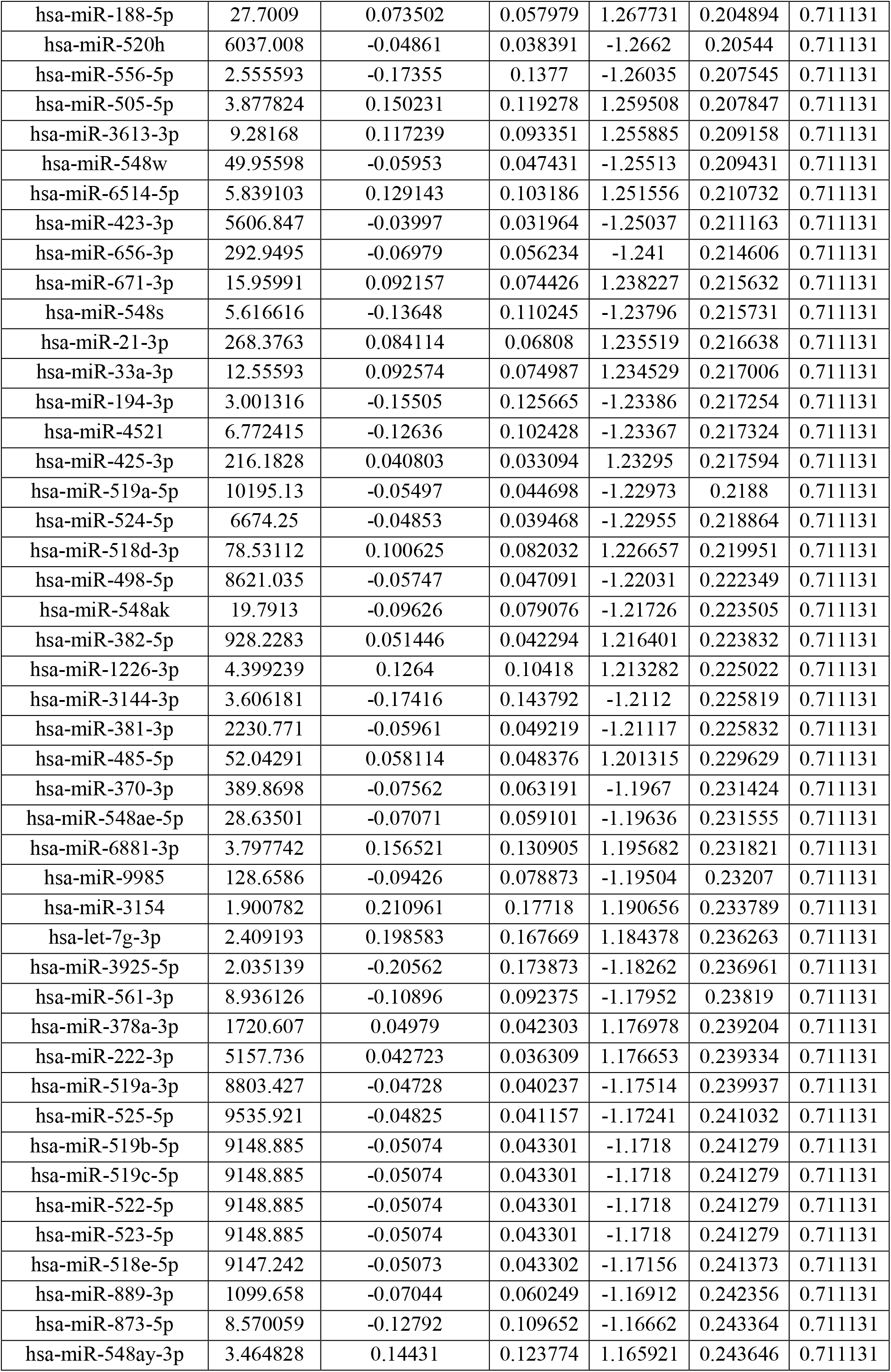
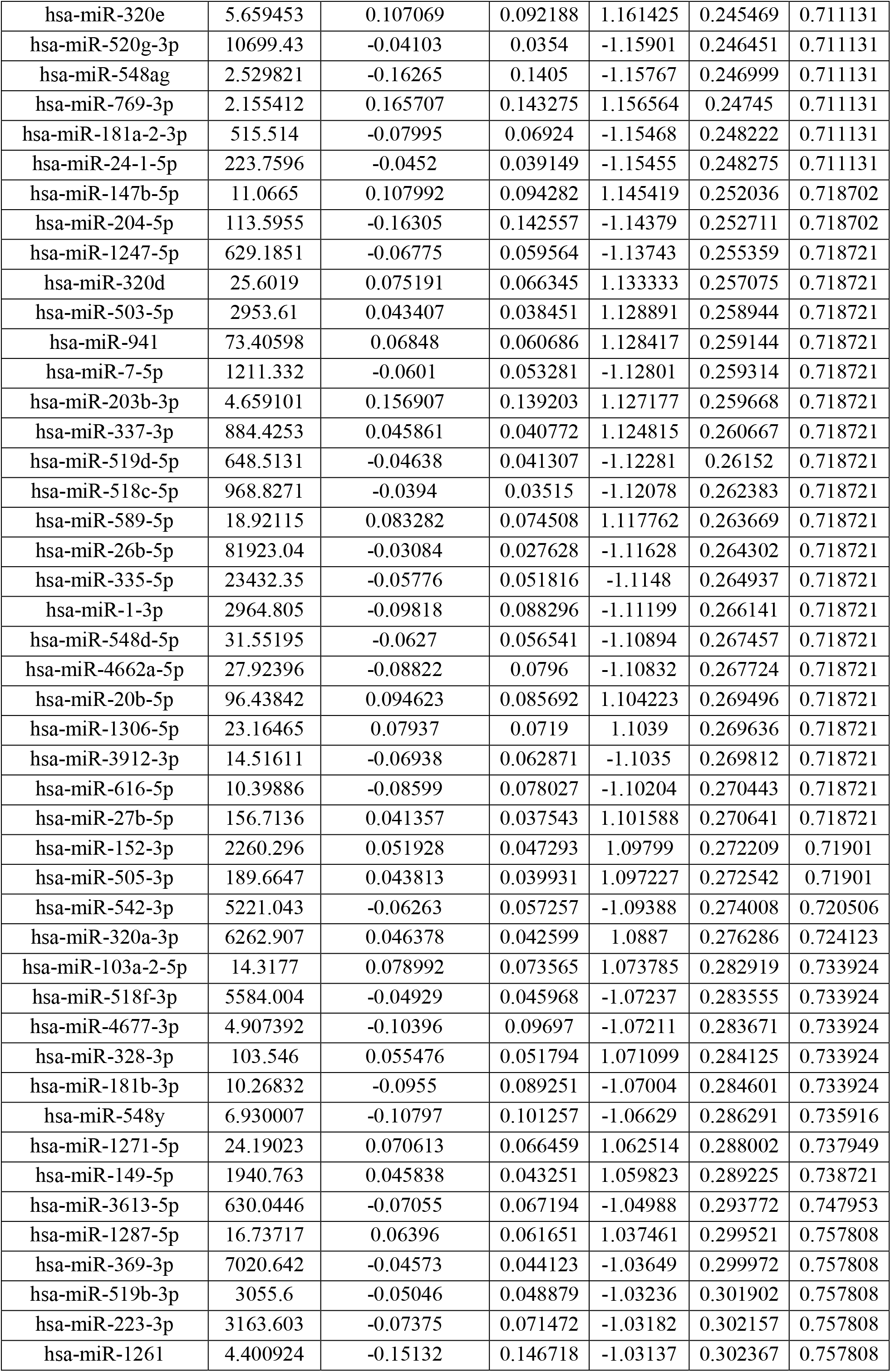
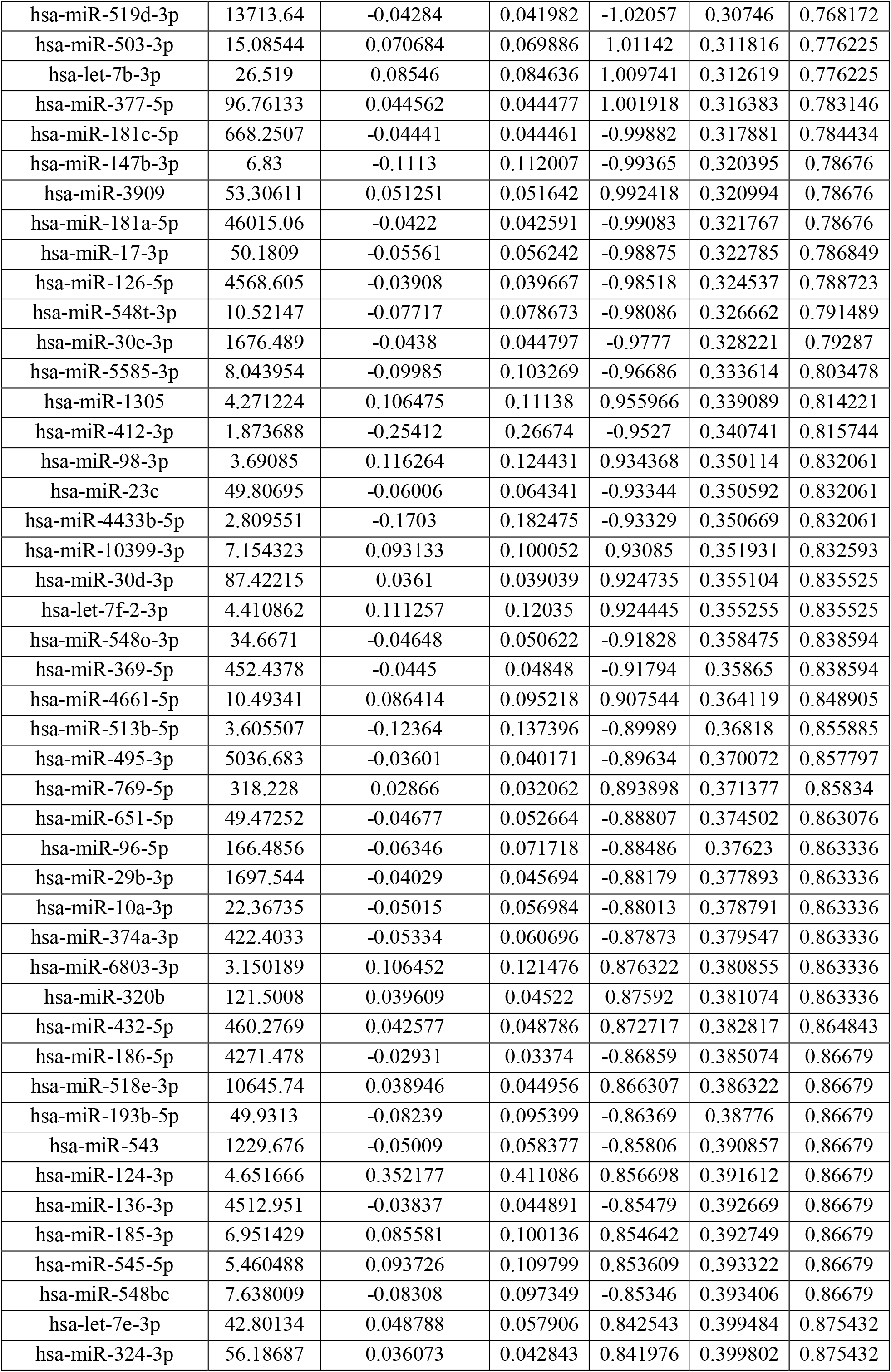
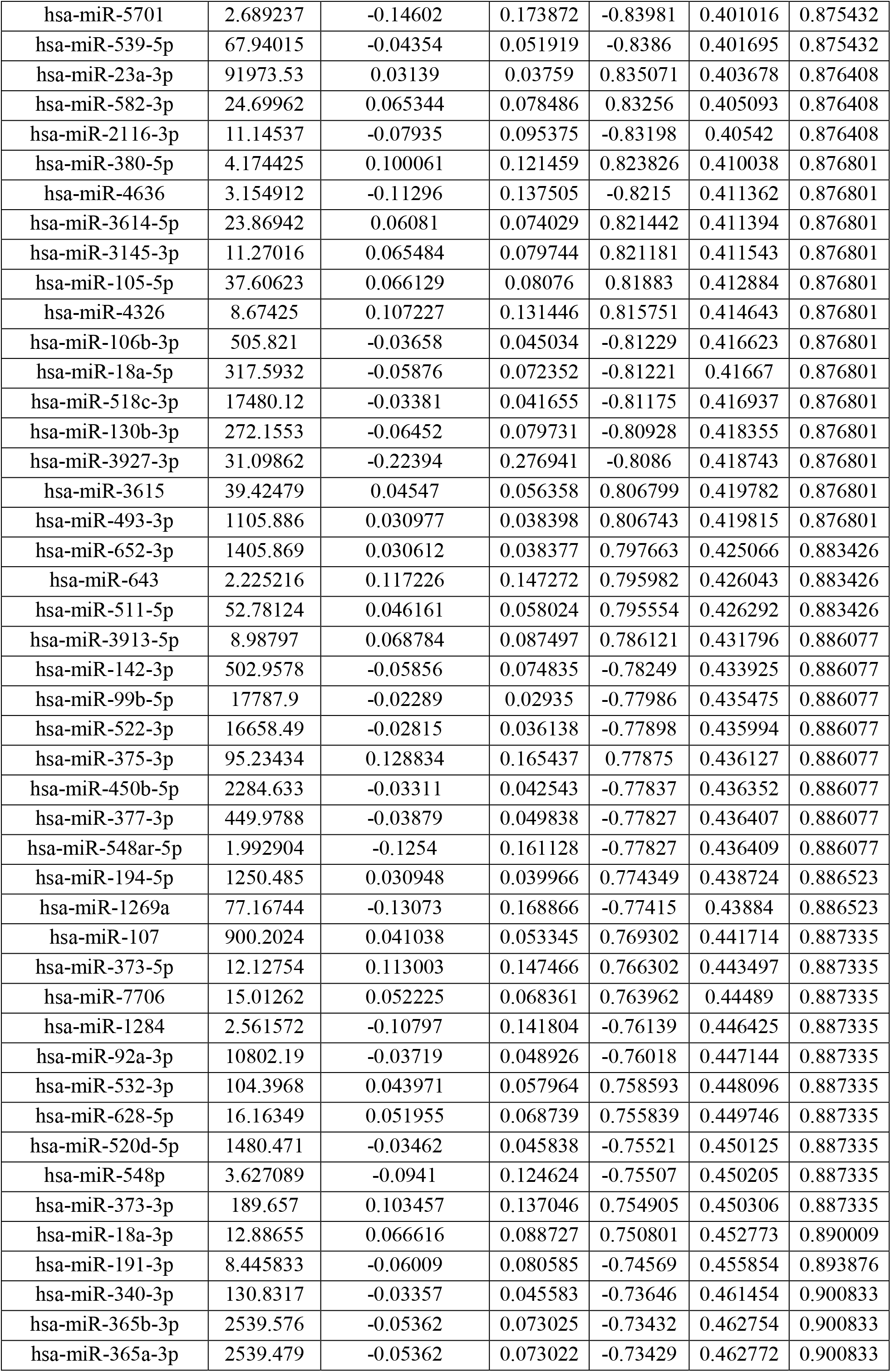
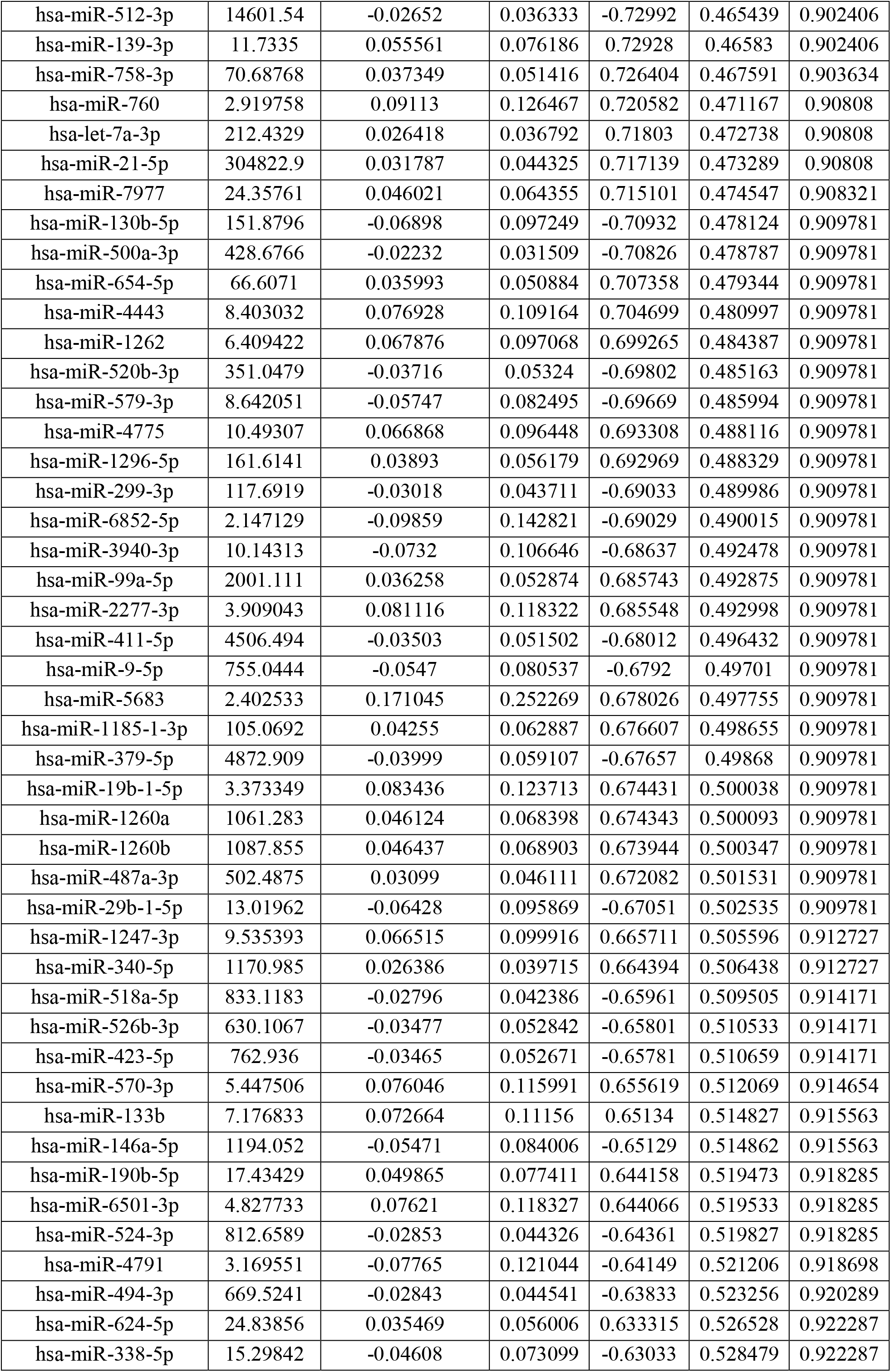
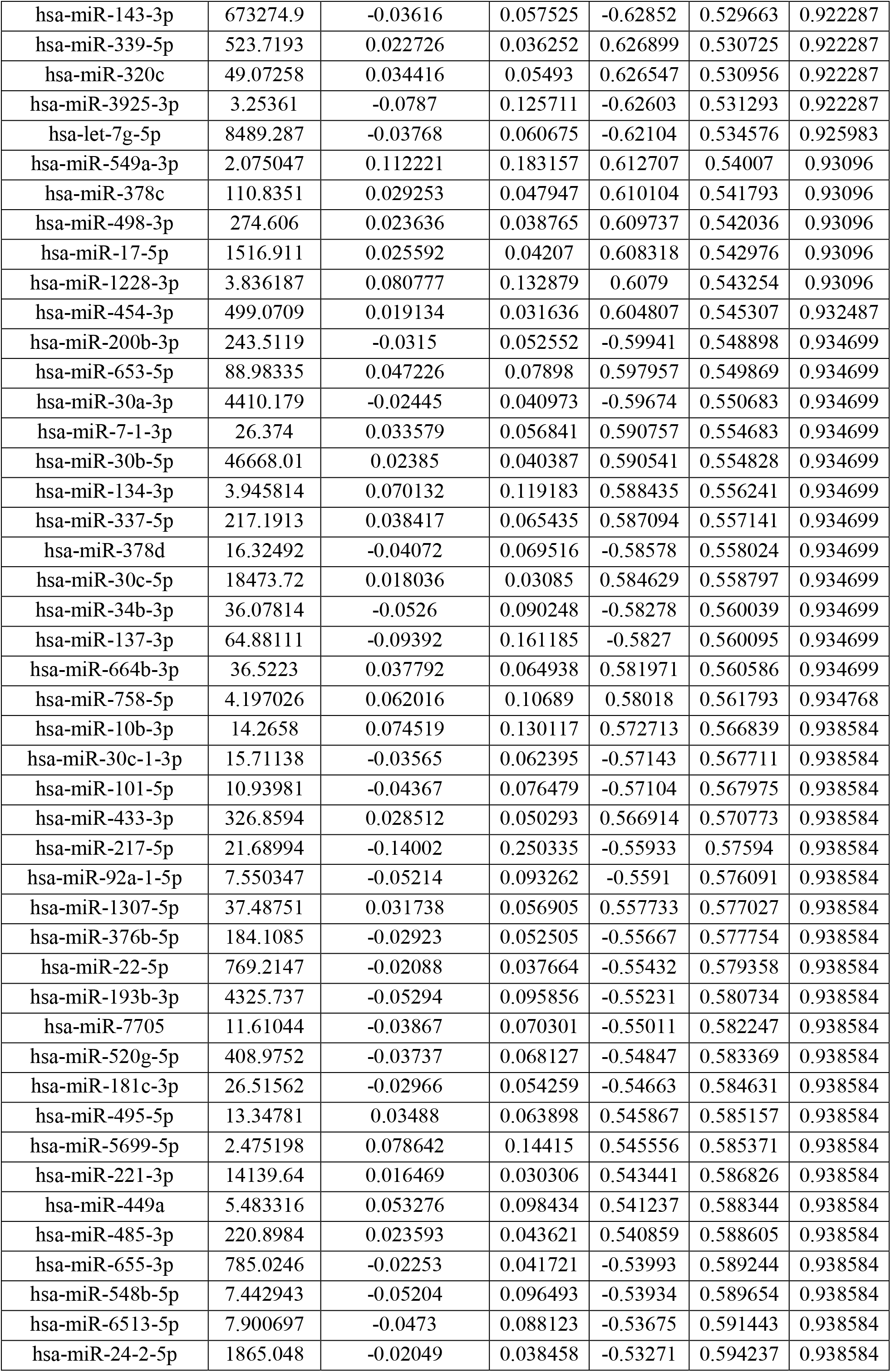
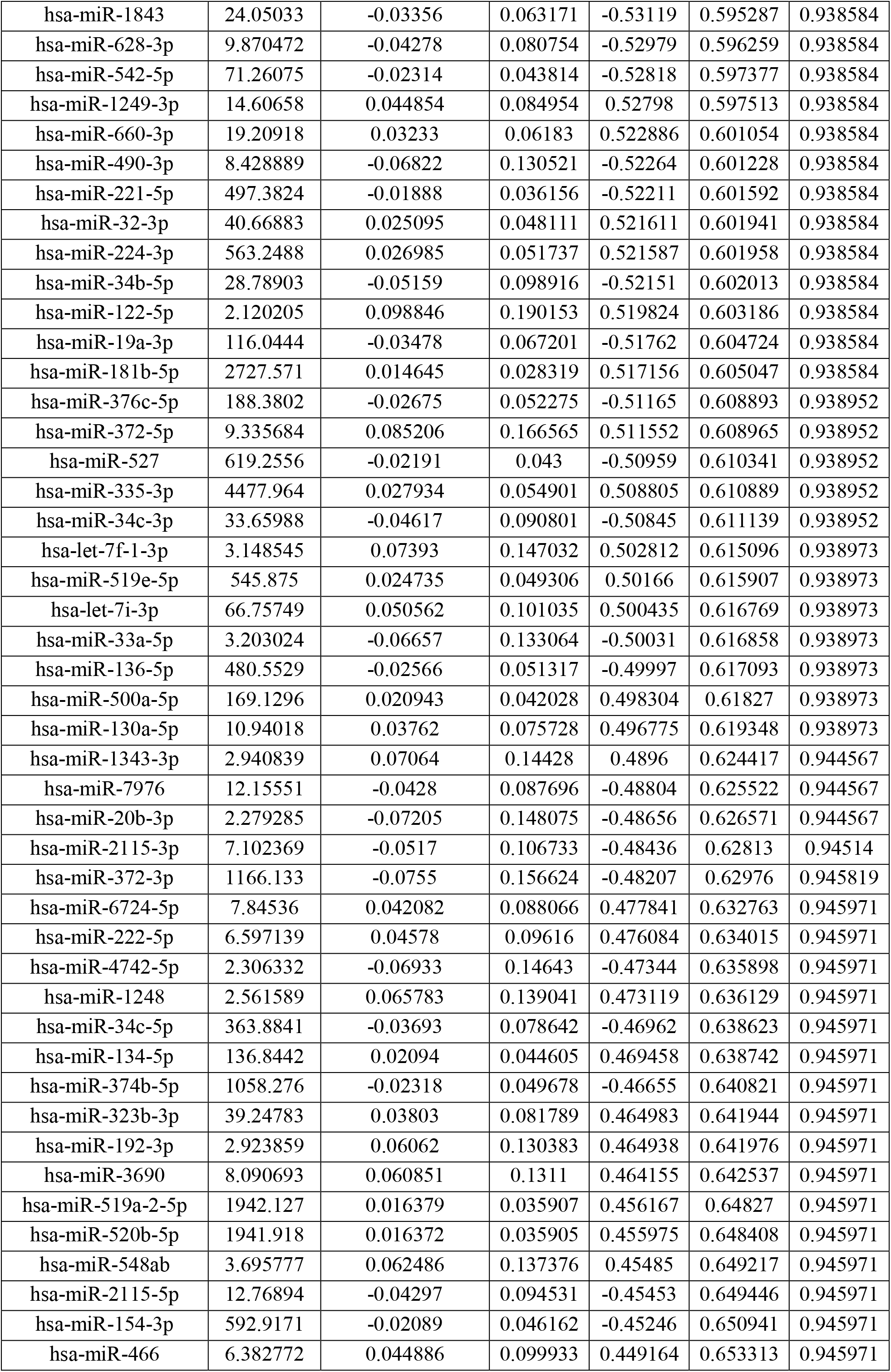
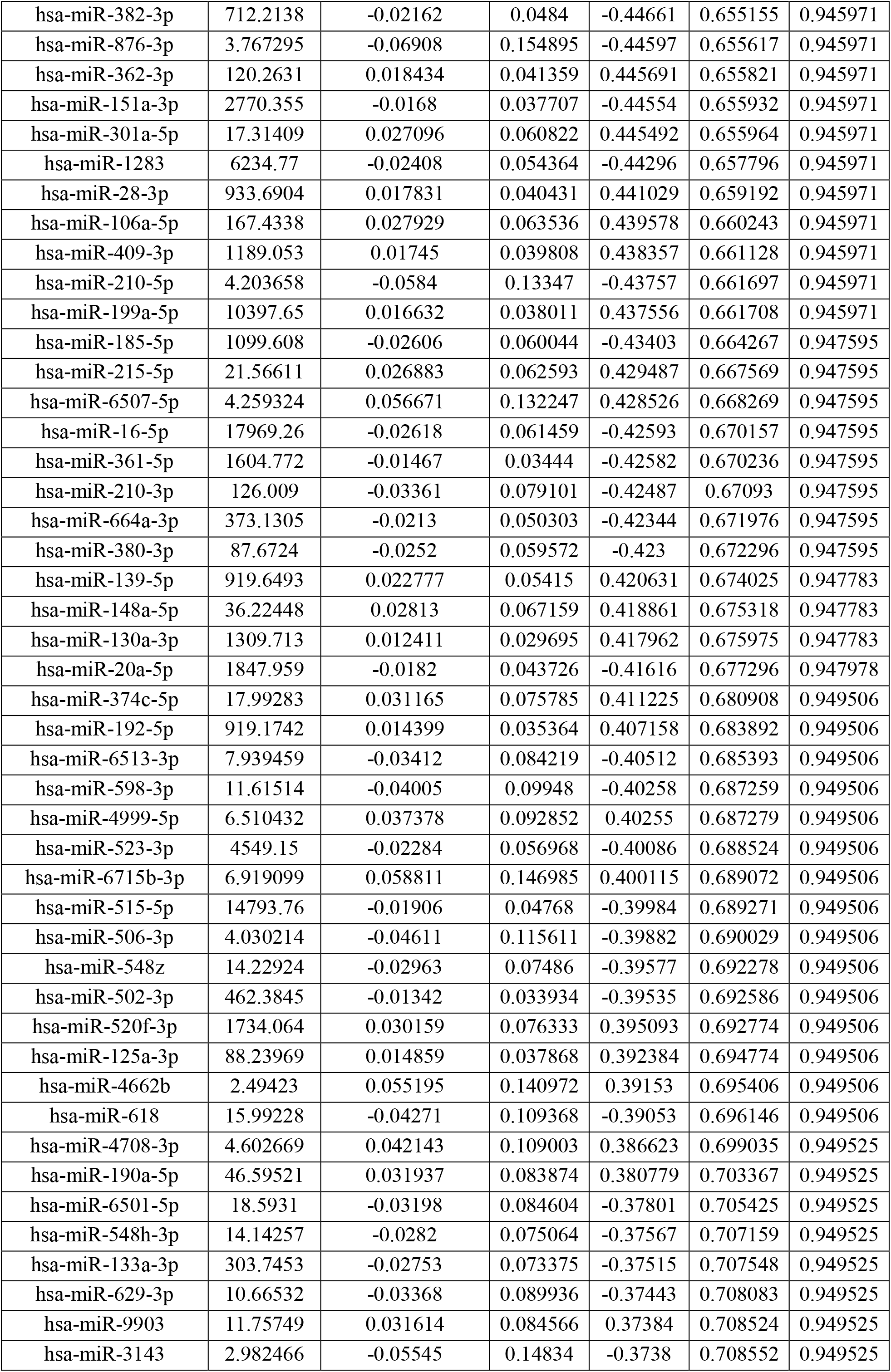
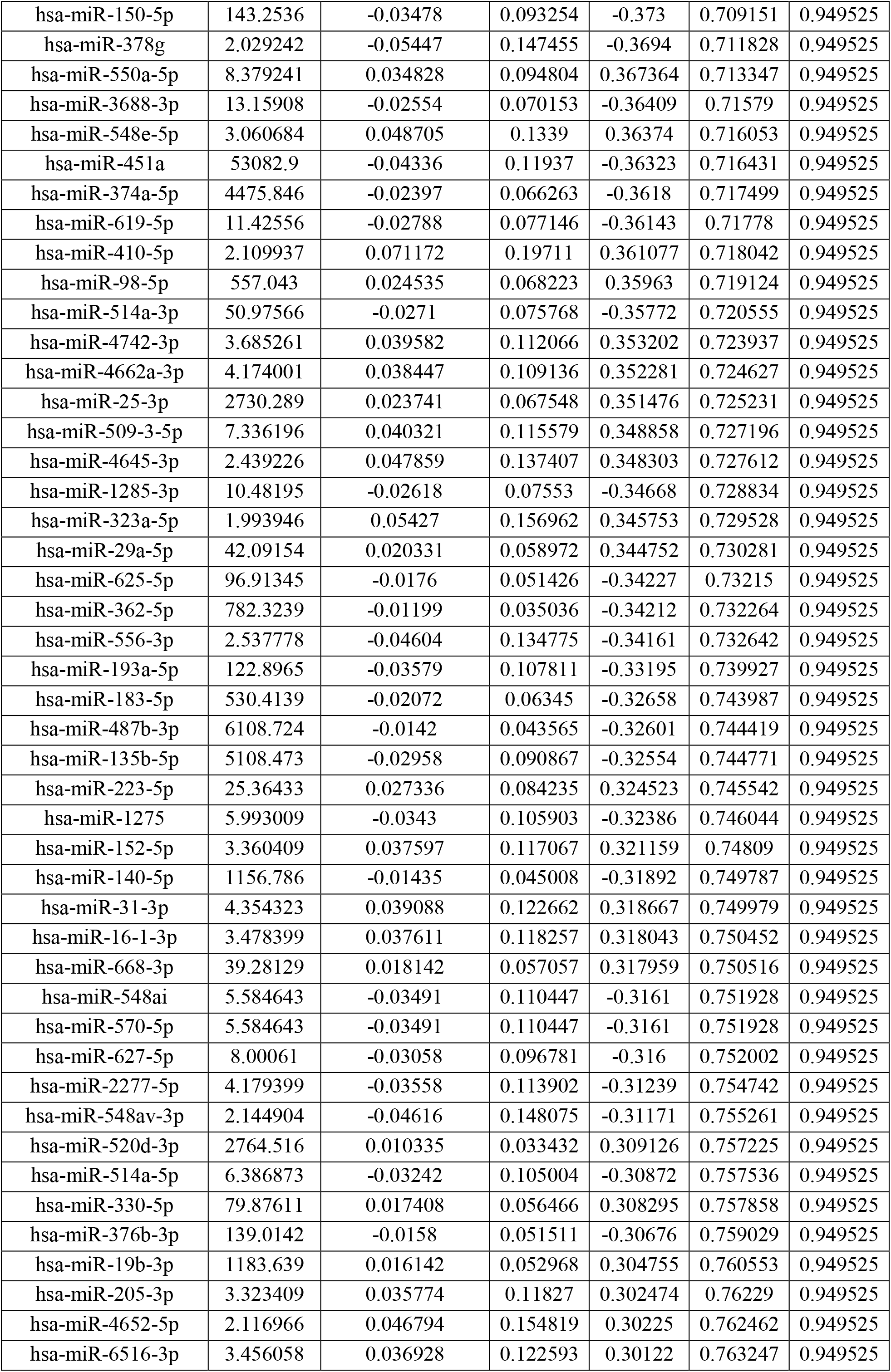
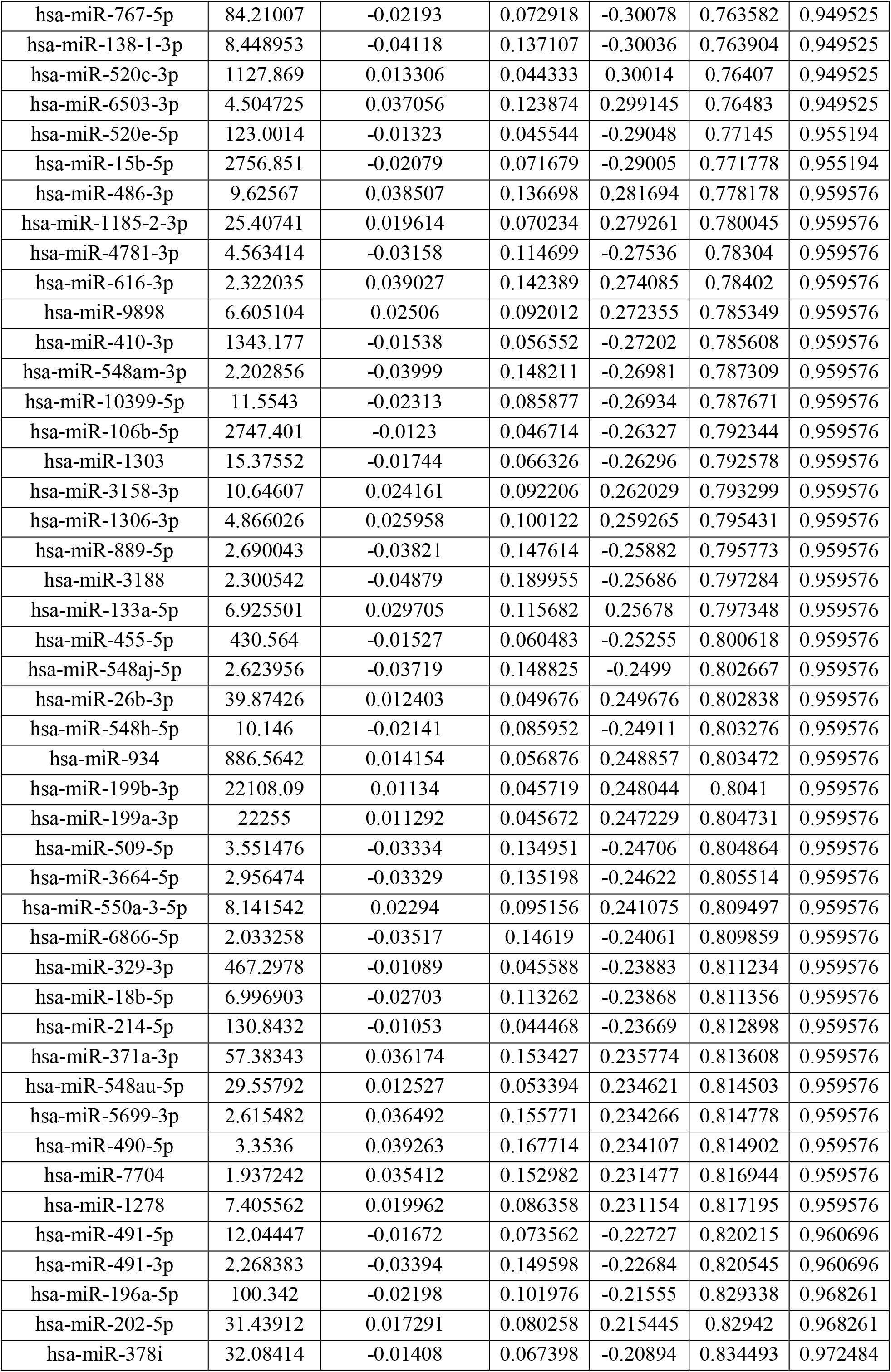
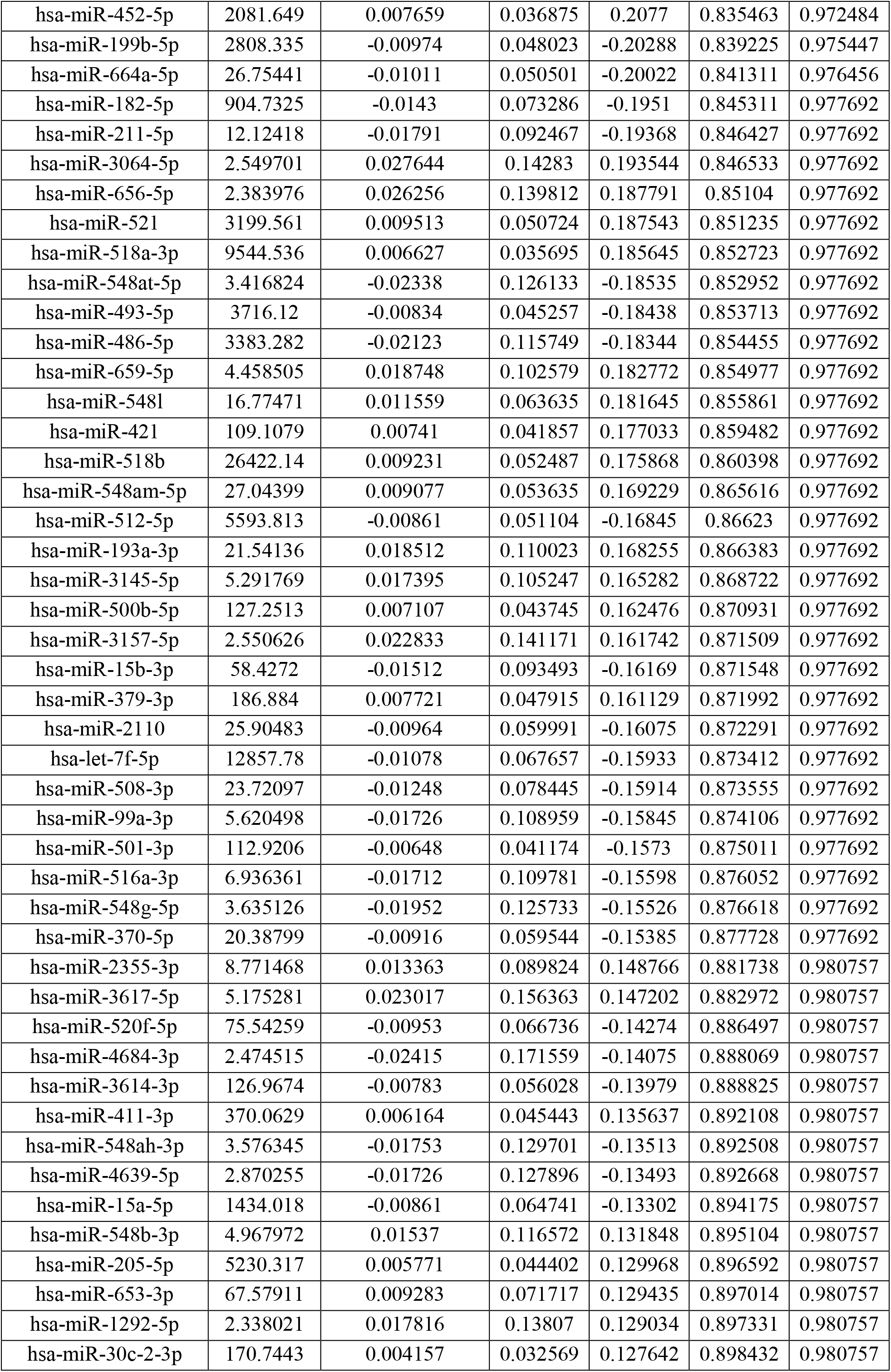
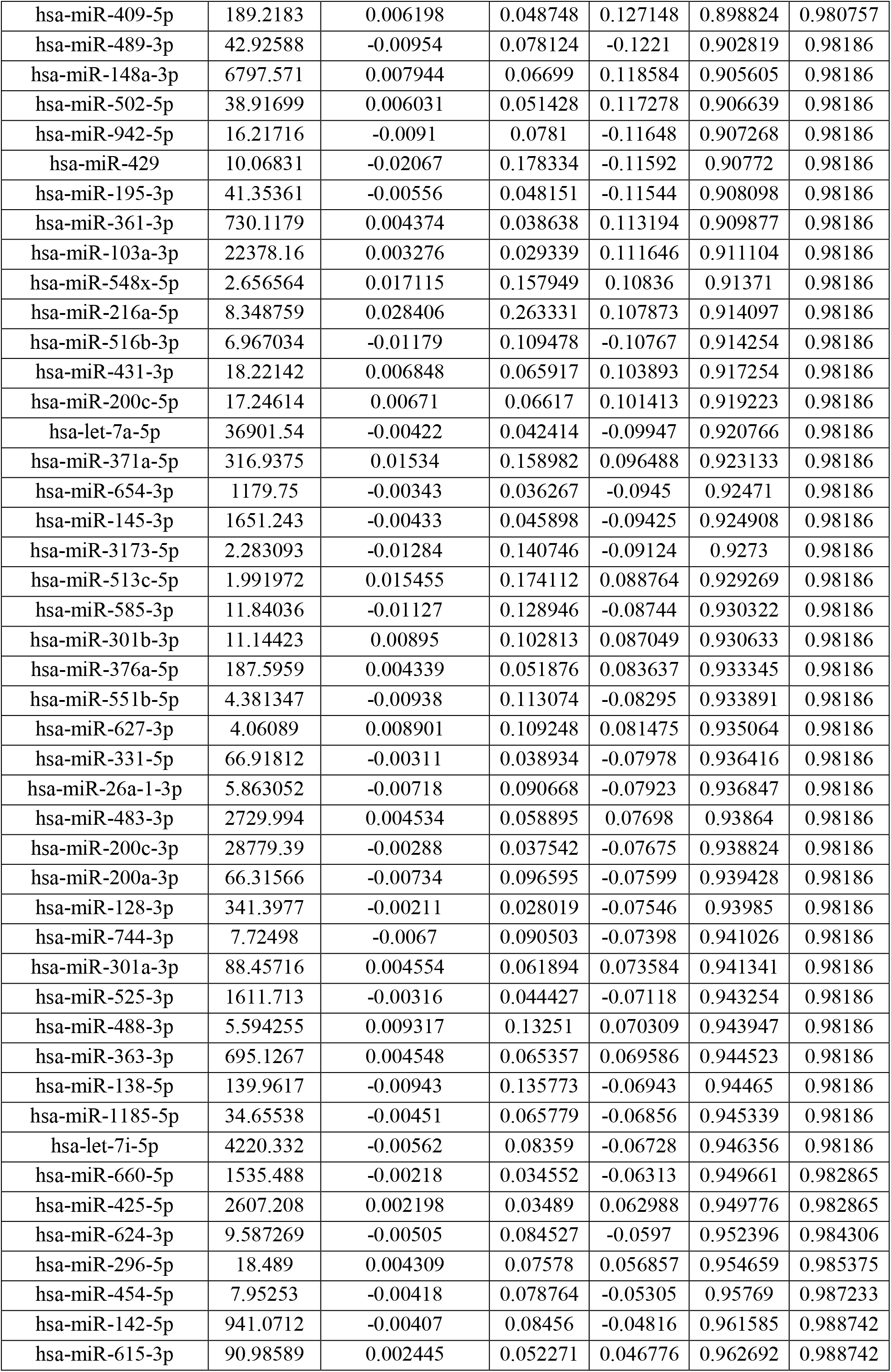
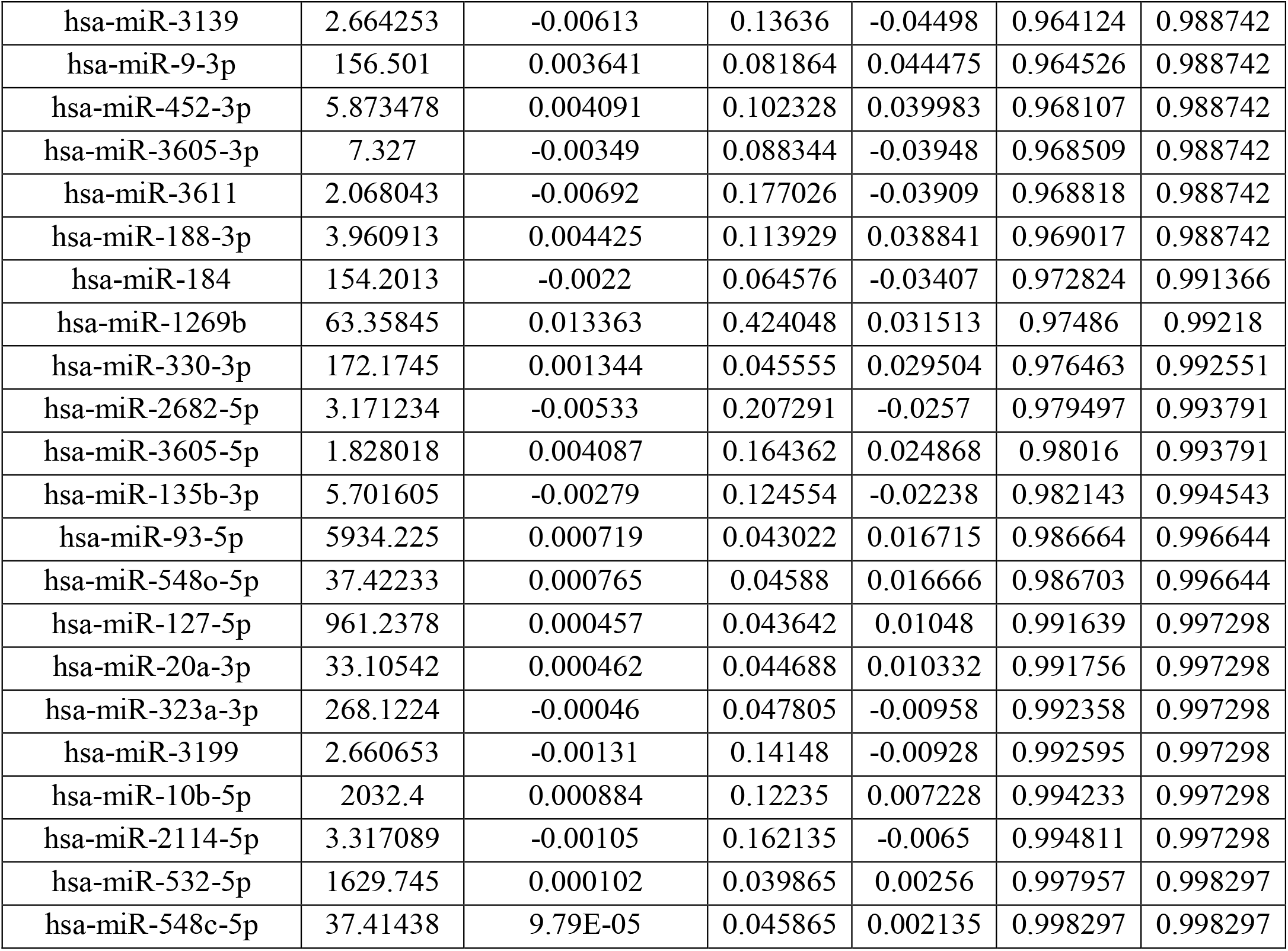
Differential Expression Analysis Results: DESeq2 Output

